# Epigenome-based prediction of gene expression across species

**DOI:** 10.1101/371146

**Authors:** Peter Ebert, Thomas Lengauer, Christoph Bock

## Abstract

**Background:** Cross-species studies of epigenetic regulation have great potential, yet most epigenome mapping has focused on human, mouse, and a small number of other model organisms. Here we explore whether existing reference epigenome collections can be leveraged for analyzing other species, by extrapolation and predictive transfer of epigenome information from established model organisms to less well annotated non-model organisms.

**Results:** We developed a methodology for cross-species mapping of epigenome data, which we used for predicting tissue-specific gene expression across twelve mammalian and one avian species. Specifically, we trained gradient boosting classifiers to predict gene expression status from reference epigenome data in human and mouse, and we applied these classifiers to epigenome profiles that were computationally transferred between species. The resulting predictions indeed identified tissue-specific differences in gene expression in the target species, thus providing initial validation of the concept of cross-species epigenome extrapolation.

**Conclusions:** Our study establishes a workflow for cross-species epigenome mapping and epigenome-based prediction of gene expression, highlighting the future potential of using epigenome maps from reference species to annotate a potentially large number of target species.

## Background

Cross-species genome analysis is widely used for dissecting evolutionary processes, identifying regulatory elements, improving genomic annotations, and studying the mechanisms underlying human diseases [1–7]. Recent progress with epigenome profiling technology has added a new dimension to genome comparisons. Large international consortia such as the International Human Epigenome Consortium [8] (which includes BLUEPRINT [9] and DEEP [10]) and the ENCODE project [11,12] produce an ever-increasing number of reference epigenomes, focused largely on human and mouse. The analysis of the resulting datasets has greatly enhanced our understanding of genomic functional elements and tissue-specific gene regulation [13–15].

Cross-species comparisons that incorporate epigenomics and functional genomics data have opened up new ways of investigating evolutionary processes, looking beyond genomic sequence conservation [16,17]. However, due to the high cost of generating epigenome data, as well as the cell-type specific and dynamic nature of epigenomic marks, current reference epigenome datasets have been limited to a handful of species, most notably human and mouse. Genome sequencing efforts for other vertebrate species rarely include epigenome profiling [18,19], which has hampered the investigation of epigenome regulation in non-model organisms and precluded systematic cross-species epigenome analyses (with a few notable exceptions [16,20,21]).

The goal of this study was to explore and evaluate cross-species extrapolation of epigenome data and epigenome-based inference of gene expression in a range of target species, based on existing reference epigenome maps for human and mouse. To that end, we established computational epigenome transfer and prediction of gene expression (Figure 1) for twelve mammalian and one avian species (Additional file 1: Figure S1). We first transferred epigenome data from our reference species (human or mouse) to the target species using whole-genome alignments. Based on the transferred epigenome data, we predicted tissue-specific gene expression in the target species using machine learning models trained and cross-validated on data from the reference species. For validation, the predictions were compared with tissue-specific expression profiles of the target species, confirming that such cross-species predictions can be useful and informative.

**Figure 1.**
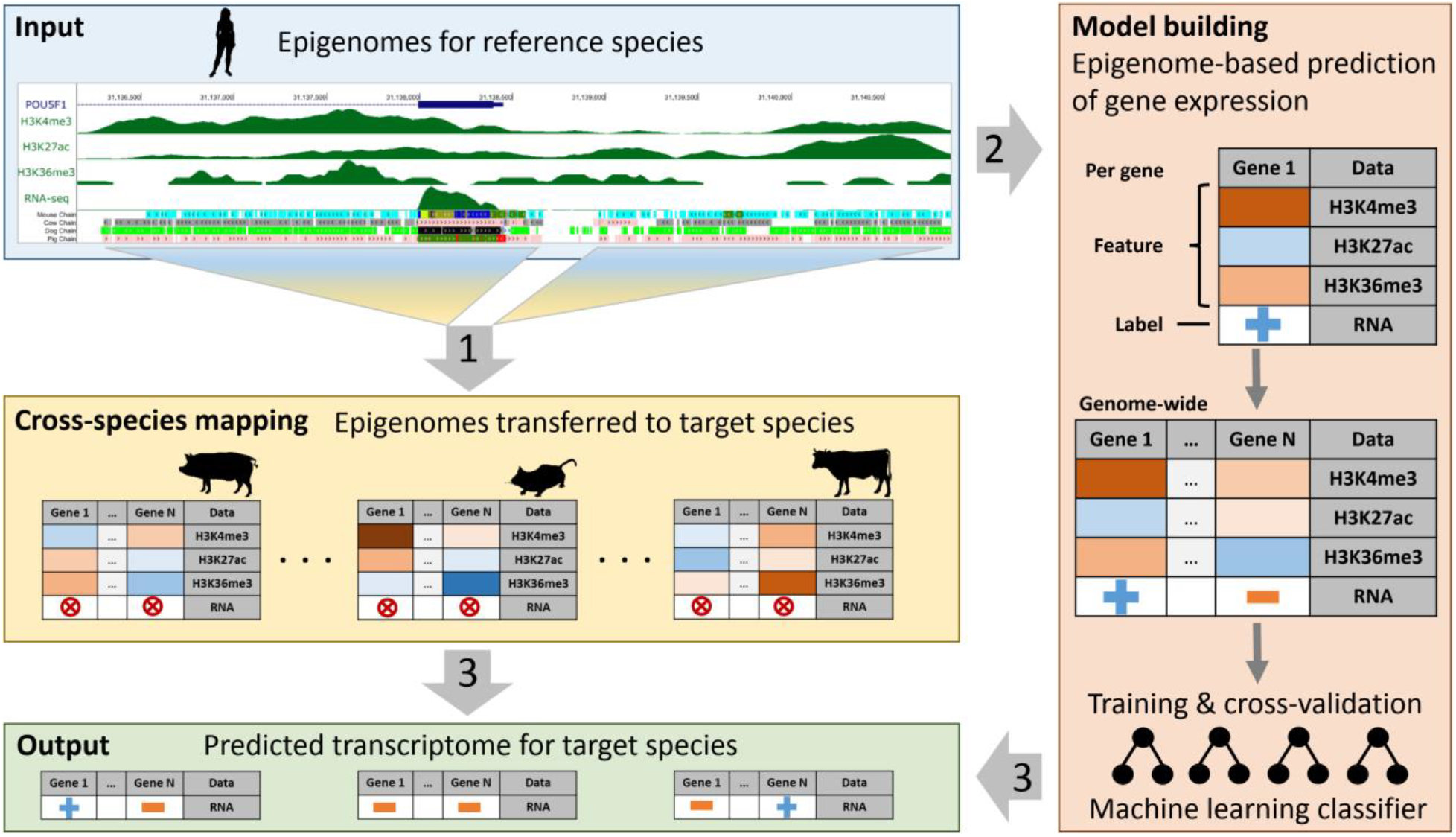
Epigenome-based prediction of gene expression across species (conceptual outline) (1) Top to middle: epigenome data from the reference species are transferred to the target species using pairwise whole-genome alignments. (2) Top to right: epigenome data in the reference species are used to train machine learning classifiers that predict gene expression status (on/high versus off/low). (3) Bottom: prediction of gene expression in the target species based on the transferred epigenome data (yellow box) and the trained machine learning classifier (red box).

## Results

### Cross-species transfer of reference epigenome data using whole-genome alignments

To test the feasibility of extrapolating and transferring epigenome data across species, we assembled a dataset comprising two reference species with extensive epigenome data (human and mouse) as well as eleven additional target species, for which we required only a reference genome and whole-genome alignments with at least one of the two reference species (Additional file 1: Figure S1). While many more vertebrate genomes have been sequenced [19,22], our species selection was based on the availability of standardized genome resources for fair and consistent performance evaluation (see Materials and Methods). The epigenome data for the reference species were obtained from public sources and include the following cell types (Additional file 2: Table S1): embryonic stem cells (human and mouse) [11,23]; CD4+ T cells (human and mouse) [9,10,24]; hepatocytes (human) [10]; whole-organ samples of liver, kidney, and heart (mouse) [23]. For each cell type, ChIP-seq profiles for three histone marks were included in the analysis: histone H3K4me3 (which marks open chromatin at gene promoters), H3K27ac (open chromatin at promoter and enhancer regions), and H3K36me3 (actively transcribed regions).

Our bioinformatics pipeline uses epigenome profiles for the reference species as input and exploits whole-genome alignments to transfer these data to conserved genomic regions in the target species. To that end, we prepared reciprocally symmetric whole-genome alignments for each pair of reference species and target species, and we used these alignments to transfer histone signal intensities. Our approach is based on the hypothesis that sequence conservation is an indicator of broader regulatory conservation within a genomic region, and that tissue-specific patterns of epigenome regulation are frequently maintained across species in sequence-conserved regions. (We tested and confirmed this hypothesis by using the transferred epigenome data to predict tissue-specific gene expression in the target species – as described further below.)

Using alignment-based epigenome transfer, we produced genome-wide, cell-type specific epigenome profiles for each target species. The non-zero part of the transferred histone signal covers a substantial amount of the overall aligned bases in the target species (ranging from human and mouse to opossum and chicken), based on the two reference species human (Figure 2) and mouse (Additional file 1: Figure S2). The distribution of the transferred histone signal followed patterns similar to that of the measured histone signal in the reference species: H3K4me3 signal was present at the smallest number of bases, consistent with this mark’s focus on gene promoters. The more widespread occurrence of H3K27ac and H3K36me3 is similarly consistent with the focus of H3K27ac on a broader set of regulatory regions (including enhancer elements) compared to H3K4me3, and H3K36me3 covers the gene body of actively transcribed genes.

**Figure 2.**
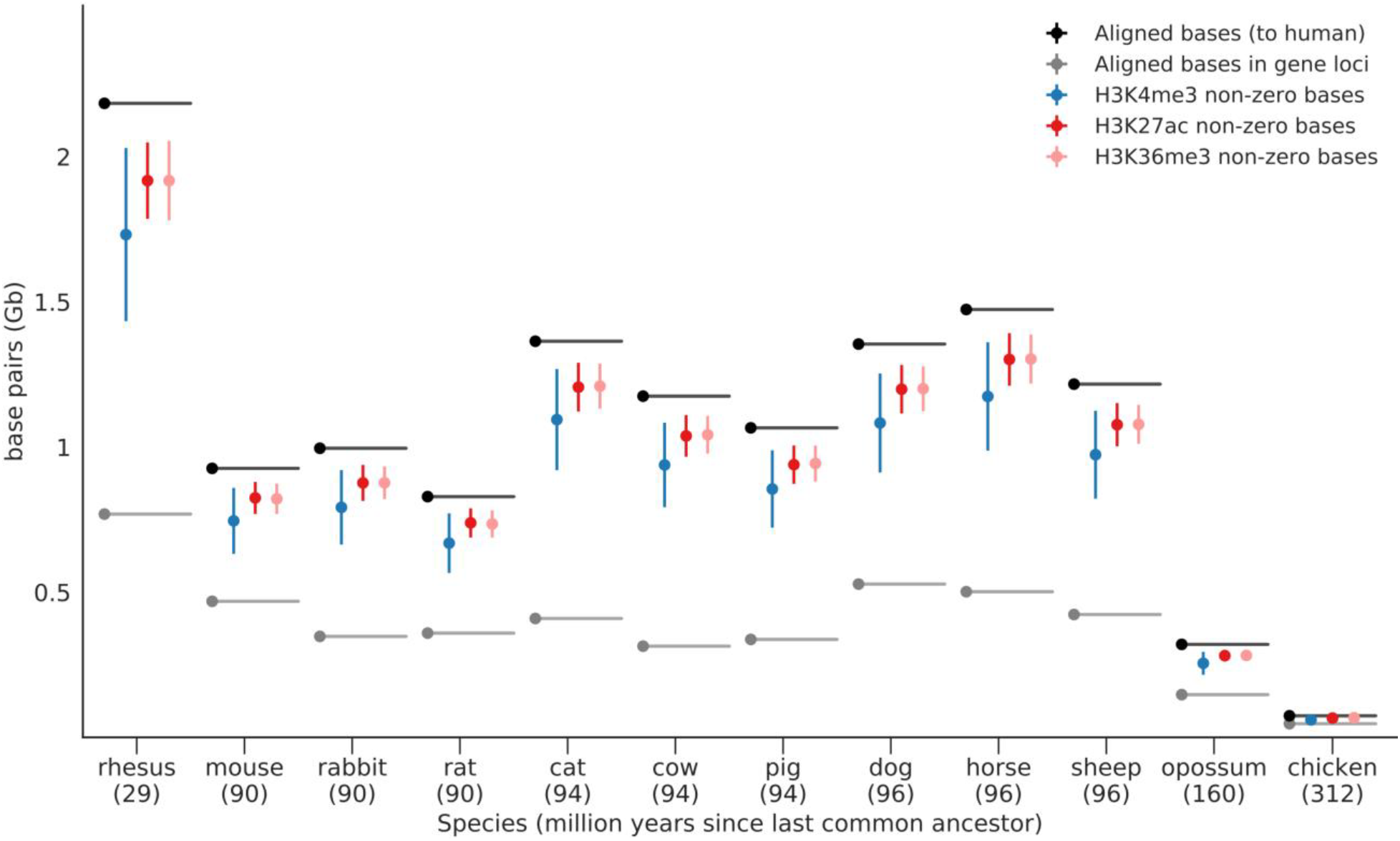
Genome-wide coverage of epigenome profiles transferred across species. Colored points indicate the average number of base pairs with non-zero signal after cross-species transfer for each histone mark (error bars denote +/- one standard deviation around the mean) between human as the reference species and twelve target species. The number of aligned bases genome-wide (black bars) and at gene loci (gene promoters and gene bodies combined, grey bars) are shown for comparison. Target species are sorted by increasing evolutionary distance to the human reference (x-axis, million years since last common ancestor indicated in parentheses).

Further analysis of average histone signal strength in gene promoters, gene bodies, and other genomic regions showed that, after cross-species transfer, the histone signal strength in the target species still resemble the distribution in the reference species (Additional file 1: Figure S3A/B): The transferred signal was generally strongest in gene bodies and in gene promoters, while it was weak outside of protein-coding genes. Since we only used the histone signal in gene promoters and in gene bodies for predictive modeling of gene expression, the low histone signal outside of these two region types is unlikely to impact the prediction analysis described below.

As a first validation for the cross-species mapping of epigenome data (Additional file 3 and 4), we examined whether the transferred epigenomes retain detectable cell-type specificity in gene promoters (for H3K4me3 and H3K27ac) and gene bodies (for H3K36me3). To that end, we focused on the comparison between human and mouse, where we have reference epigenome data for both species, and we found that correlations between transferred and measured epigenomes were usually highest when the cell type was matched (Figure 3). For example, the comparison between transferred and measured epigenomes in mouse resulted in a Pearson correlation of 0.75 for H3K4me3, 0.61 for H3K27ac, and 0.71 for H3K36me3 in CD4+ T cells.

**Figure 3.**
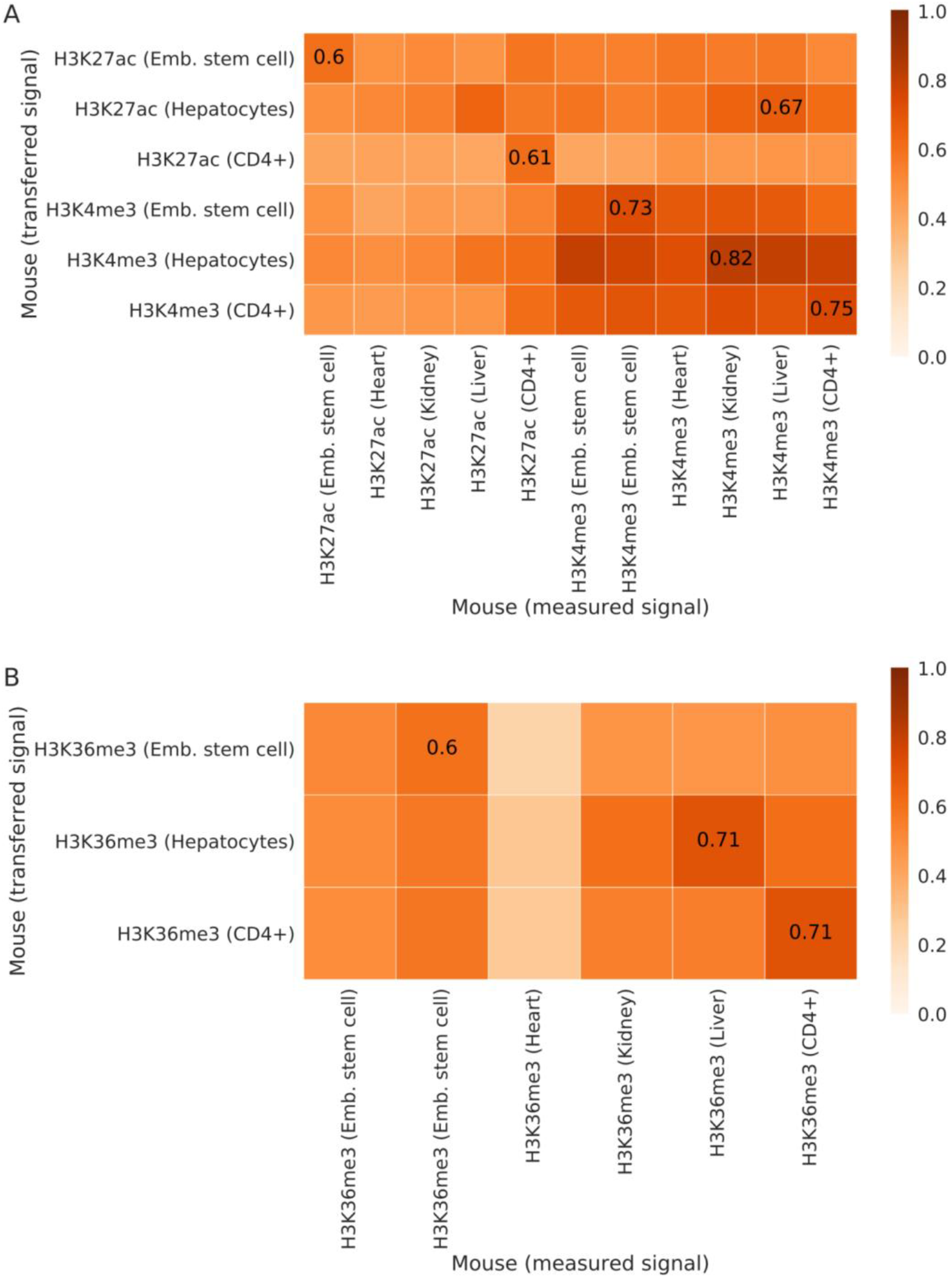
Tissue-specific correlation of measured and transferred epigenomes. Heatmaps showing pairwise Pearson correlations of measured and transferred histone signals for H3K4me3 and H3K27ac at gene promoters (panel A) and for H3K36me3 in gene bodies (panel B), using human as reference species and mouse as target species.

To substantiate these findings, we performed region set enrichment using the LOLA software [25] on the top 5% gene promoters with the strongest transferred signal for H3K4me3, and on the top 5% gene bodies with strongest signal for H3K36me3. We consistently observed cell type specific enrichments (Addition file 5), as illustrated by the LOLA enrichments for CD4+ T cells using mouse as reference and human as target species (Figure 4A/B) and by the LOLA enrichments for liver/hepatocytes using human as reference and mouse as target species (Figure 4C/D). In both cases, the LOLA results showed the expected enrichment for tissue-specific histone marks associated with active promoters (H3K4me3) and actively transcribed genes (H3K36me3), respectively. The annotated cell type of the enriched region sets was the same as (or closely related to) the cell type of the transferred epigenomes, supporting that our cross-species epigenome transfer method retained the characteristic cell-type specificity of epigenome data.

**Figure 4.**
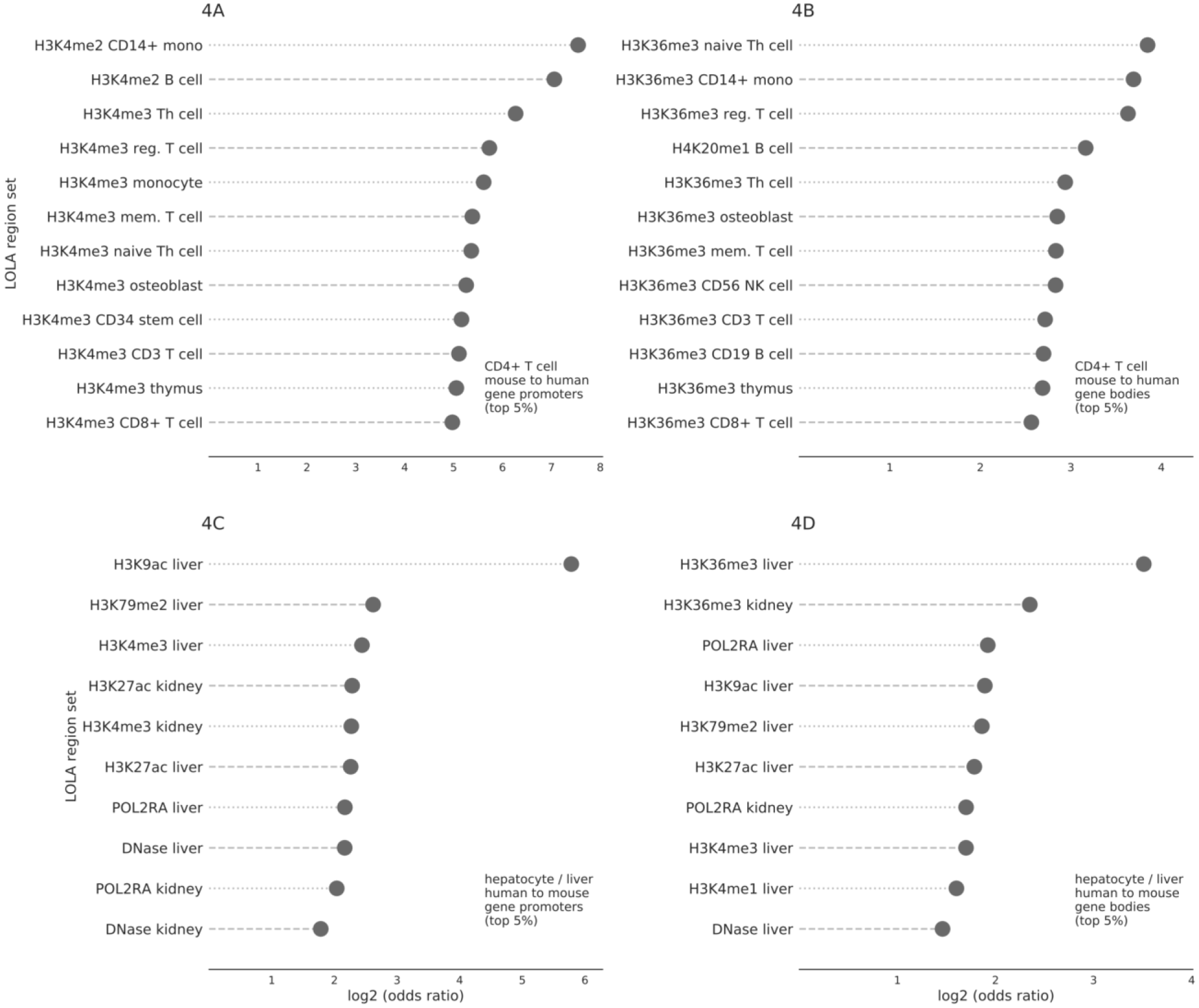
Genomic region enrichment analysis for transferred epigenomes. Selected results of LOLA analyses for gene promoters (panel A and C) and gene bodies (panel B and D) that were ranked among the top 5% based on transferred epigenome signal intensities for H3K4me3 (gene promoters) or H3K36me3 (gene bodies). Top panels show region set enrichment for CD4+ T cell epigenomes transferred from mouse to human (panel A and B); and bottom panels show region set enrichment for hepatocyte (liver) epigenomes transferred from human to mouse (panel C and D). Effect size (log2 of the odds ratio) is indicated on the x-axis. The false discovery rate estimate (q-value) is smaller than 10e-12 in all cases. Complete results of the LOLA analysis are shown in Additional file 5.

### Prediction of gene expression using epigenome data transferred across species

To assess the biological information carried by the transferred epigenomes in a larger number of species (namely in those for which no suitable epigenome data are available for validation), we tested whether we could predict gene expression in the target species based on the transferred epigenomes. It is well-established that gene expression can be predicted from epigenome data [26–28], and we would expect to observe better-than-random accuracies predicting gene expression based on transferred epigenome profiles if these profiles indeed captured relevant regulatory biology. Moreover, to make the validation more stringent, we can exploit the cell-type specific character of epigenome data and try to predict cell-type specific patterns of gene expression.

As our experimental reference dataset against which we evaluated the epigenome-based predictions, we obtained transcriptome data from various public sources covering a range of species and cell types: embryonic stem cells for human and mouse [11,12]; CD4+ T cells for human and mouse [24,29]; hepatocytes for human [10]; whole-organ samples of liver, kidney and heart for all species except human [30–32]; and a blood sample for opossum (Additional file 2). In this transcriptome dataset, we observed consistent clustering by cell type rather than by species, both for 1-to-1 gene orthologs [33] between each pair of species (Figure 5) and for those genes that were conserved across all 13 species (Additional file 1: Figure S4).

**Figure 5.**
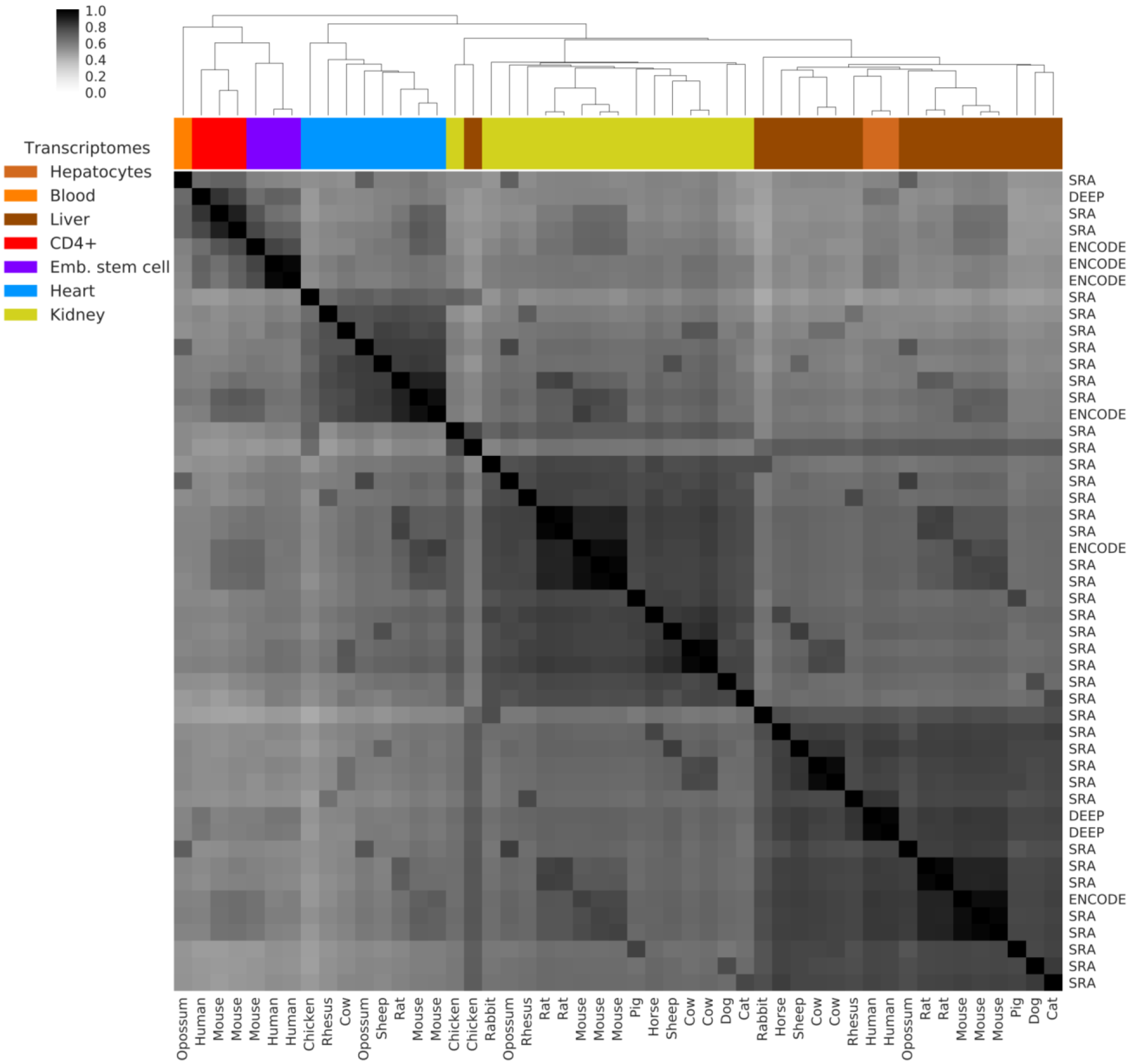
Tissue-specific clustering of the cross-species transcriptome dataset. Hierarchical clustering of pairwise Pearson correlation coefficients between transcriptome profiles of 13 species used for gene expression prediction and model validation. Correlations were calculated across all 1-to-1 orthologous genes for each species pair. Color bars at the top indicate tissue of origin. Columns are labeled with species of origin, and row labels indicate data source (project name or “SRA” for data downloaded from SRA/ENA; see Additional file 2 for details).

We implemented a machine learning approach to predict gene expression profiles in the target species based on epigenome data in the reference species (see Figure 1 for an overview). Using only data from the reference species, binary classifiers were trained to predict gene expression as either on/high or off/low. The prediction attributes were obtained by averaging histone signals for H3K4me3/H3K27ac in gene promoters and for H3K36me3 within the gene body, restricted to those subregions that were covered by the cross-species alignment used for epigenome transfer. A threshold of one transcript per million (1 TPM) was applied to label genes as on/high or off/low. We observed consistently high prediction performance for the two reference species, with crossvalidated, test-set only, receiver operating characteristic (ROC) area under curve (AUC) values of 0.89 (human to mouse) and 0.90 (mouse to human), resulting in a sensitivity of 78% (human) and 76% (mouse) at a specificity of 90% (Figure 6A/B).

**Figure 6.**
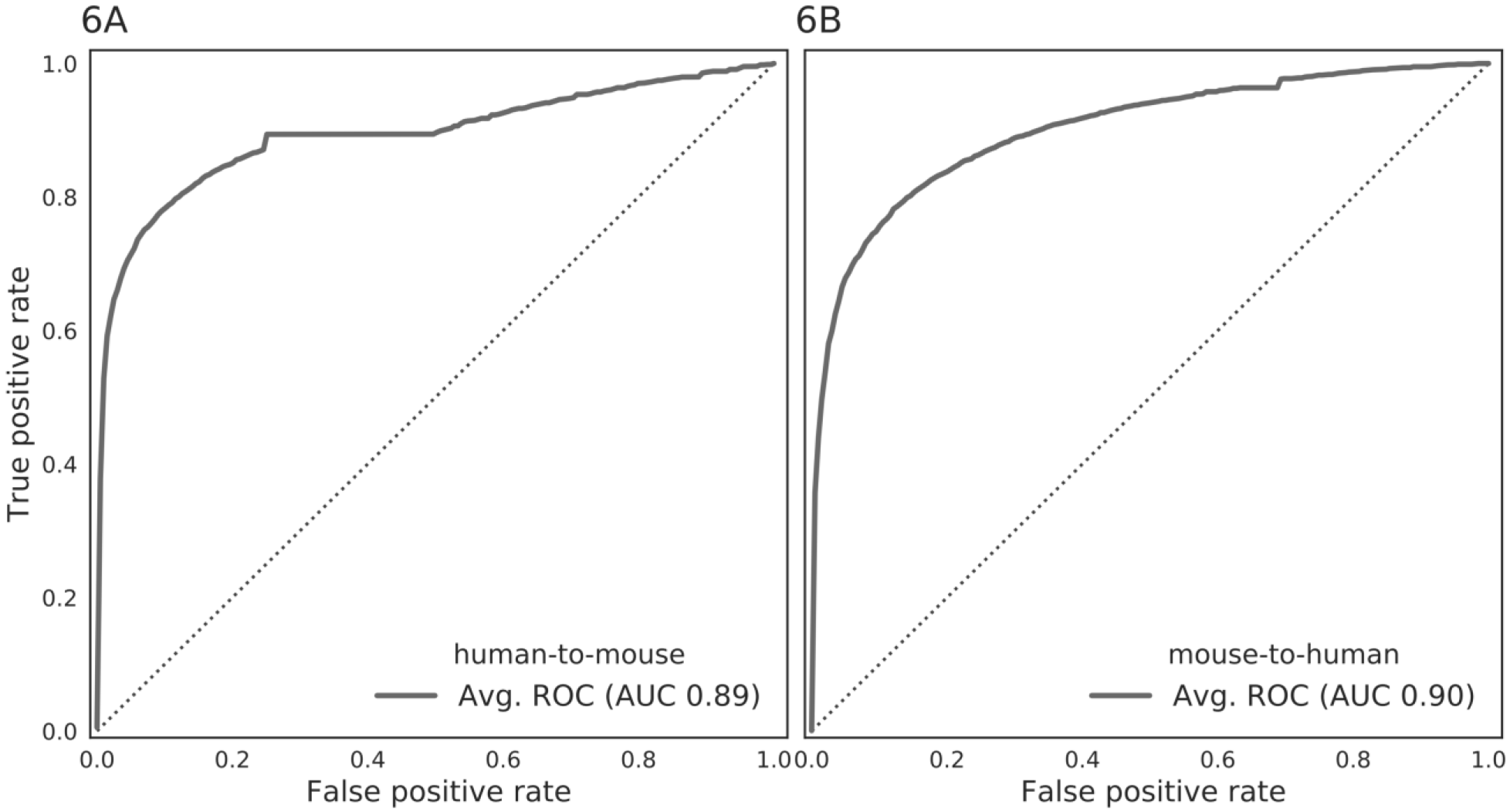
Performance for predicting gene expression from transferred epigenomes in human/mouse. ROC curves showing the cross-validated test-set performance predicting gene expression status in mouse based on transferred epigenome data from human (panel A) and vice versa (panel B), averaged across all included samples and cell types in both the reference and the target species. The dashed diagonal line represents the expected performance of a random classifier (AUC = 0.5).

Having validated the epigenome-based classifiers in the reference species, we next applied them to all target species, using the transferred epigenome data as input. First, we predicted gene expression in each target species independent of cell type, averaging across transferred epigenome profiles and transcriptomes in each target species. We observed high prediction accuracies for all target species, with ROC AUC values ranging from 0.87 to 0.81 when using human as reference (Figure 7A), and from 0.89 to 0.83 when using mouse as reference (Figure 7B). The average sensitivity was 67% (human reference) and 73% (mouse reference) at a specificity of 90%.

**Figure 7.**
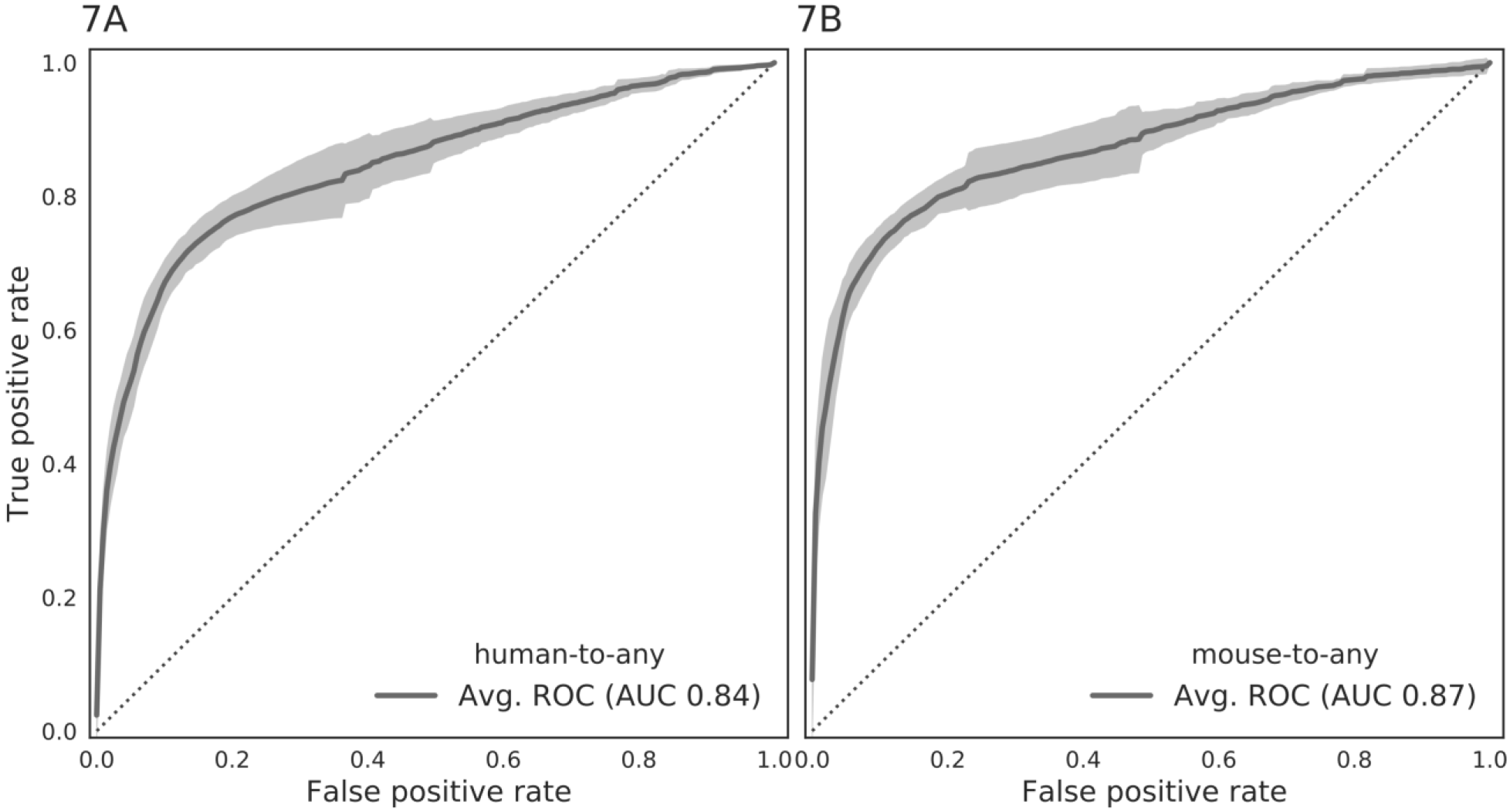
Performance for predicting gene expression from transferred epigenomes in all species. ROC curves showing the cross-validated test-set performance predicting gene expression status based on transferred epigenomes from human (panel A) or mouse (panel B) as reference species and the remaining twelve species as target species. Epigenomes and transcriptomes were averaged across all included samples and cell types in both the reference and the target species. Shaded areas represents +/- 1 standard deviation around the mean ROC curve. The dashed diagonal line represents the expected performance of a random classifier (AUC = 0.5).

As an additional evaluation, we tested to what degree our gene expression predictions were celltype specific, and whether they indeed showed the highest accuracy for the matched cell type in the target species. When comparing AUC values for tissue-matched prediction to those obtained by cross-tissue prediction, the tissue-specific predictions consistently outperformed the tissue-agnostic predictions across all investigated target species and for both human (Figure 8A) and mouse (Figure 8B) as our reference species. Aggregating AUC values across species, the difference between the mean AUC values for the tissue-matched tests and the cross-tissue controls was highly statistically significant (one-sided Mann-Whitney-U, p-value < 10e-9).

**Figure 8.**
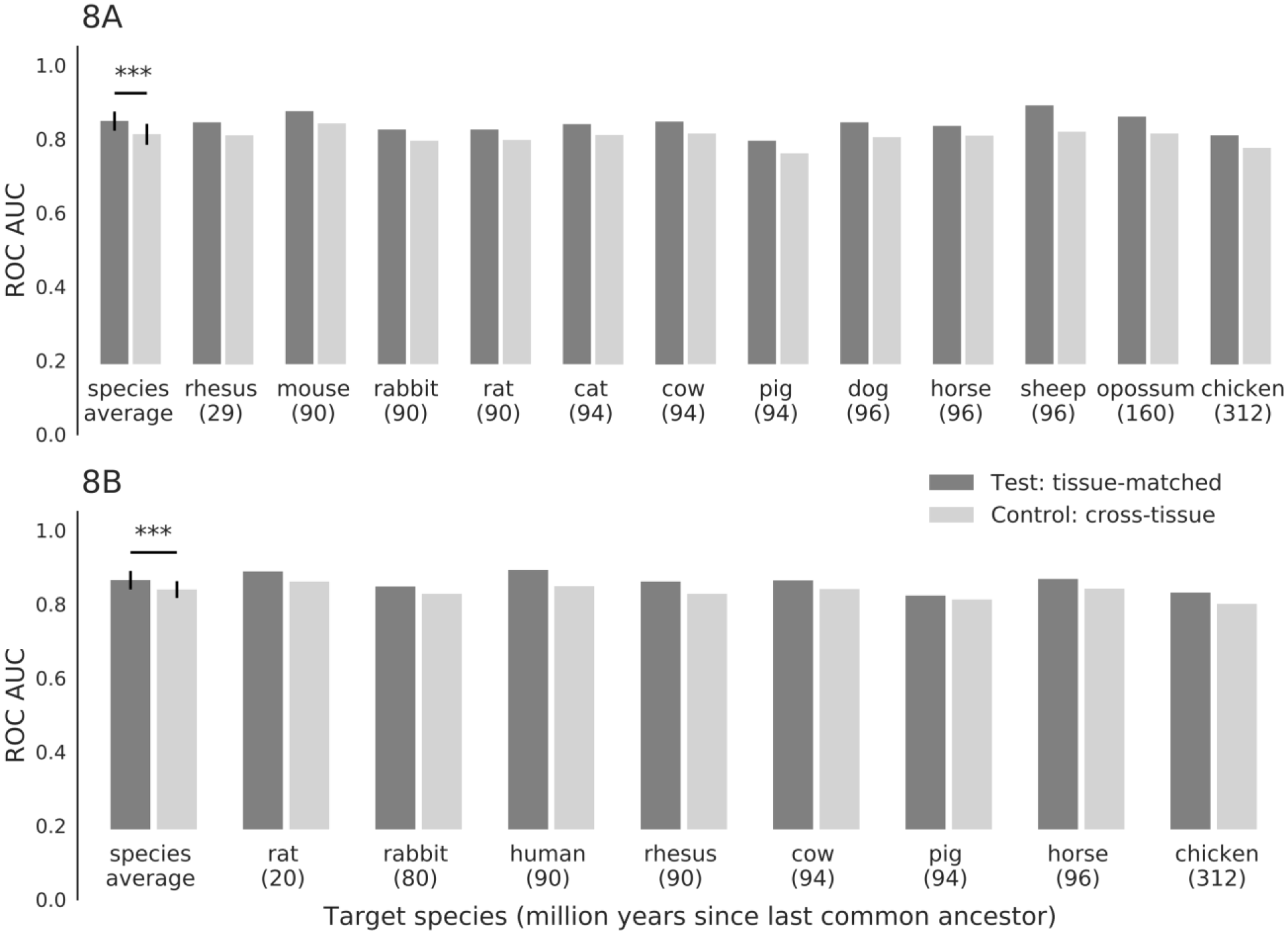
Performance gain of tissue-specific prediction of gene expression from transferred epigenomes. ROC AUC values (calculated as in Figure 7) comparing the performance of gene expression prediction based on tissue-specific (dark grey) as well as average (light grey) transferred epigenomes, using human (panel A) or mouse (panel B) as reference species and the remaining twelve species as target species. The first bar shows the aggregation of mean AUC values across all species. The difference in mean AUC for these aggregated values is statistically significant for both reference species (one-sided Mann-Whitney-U, *** p-value < 10e-9).

### Comparison of epigenome-based and orthology-based prediction of tissue-specific expression

To benchmark our epigenome-based predictions of gene expression, we compared their performance to that of an alternative (and complementary) approach that is based on gene orthology. Specifically, for all 1-to-1 gene orthologs between each pair of species, the orthology-based method predicts a gene to be expressed in a certain cell type of the target species if – and only if – it is expressed in the corresponding cell type of the reference species. This method can predict expression only for genes that have an annotated 1-to-1 ortholog in the reference species, which limits its scope and applicability to a subset of genes (Annotation file 2: Figure S5).

For a systematic comparison between the epigenome-based and orthology-based methods, we evaluated their performance on various subsets of genes selected using the following approach: we ordered all genes in the target species by increasing levels of DNA sequence conservation in the gene body, and we used all genes above a certain threshold as our evaluation set. We then plotted the performance gain of the epigenome-based method (calculated as the surplus of correct predictions over the orthology-based method across all target species) for human and mouse as our reference. In this analysis, the epigenome-based method resulted in approx. 20% more correct predictions than the orthology-based method (Figure 9A).

**Figure 9.**
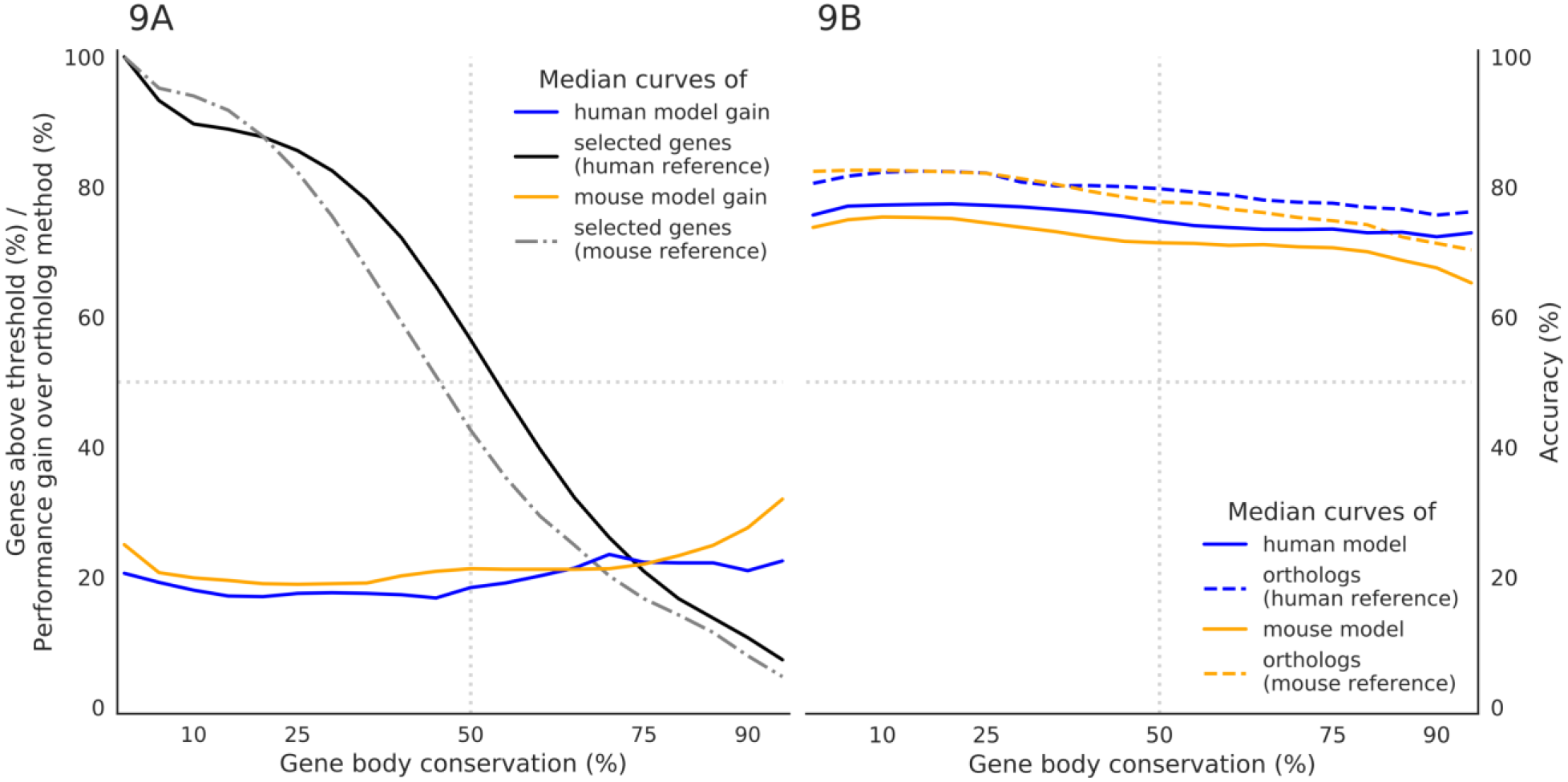
Comparison of gene expression prediction based on transferred epigenomes vs. gene orthology. (A) Performance gain of epigenome-based prediction of gene expression over the orthology-based approach using human (blue) or mouse (orange) as reference species, plotted for different thresholds on the gene body conservation (x-axis). The number of selected genes for each threshold is also indicated (black and grey lines). Performance is measured as number of correct predictions (true positives and true negatives), and the surplus of correct predictions of the epigenome-based prediction models over the orthology-based approach is transformed into a percentage value that is comparable across species. (B) Prediction accuracy of epigenome-based (solid lines) and orthology-based (dashed lines) prediction of gene expression (solid lines) and the corresponding orthology-based predictions (dashed lines) using human (blue) or mouse (orange) as reference species, plotted for different thresholds on the gene body conservation (x-axis). All curves (panel A and B) represent median values of all gene expression predictions aggregated per reference species over all target species and over all cell types. Genes in the target species are sorted by increasing gene body conservation along the x-axis.

Both methods displayed stable prediction accuracy over a broad range of gene conservation values (Figure 9B). While the performance gain of epigenome-based prediction was higher for stringent thresholds on gene body conservation (Figure 9A), only relatively few genes pass these stringent thresholds, and a more inclusive threshold may increase the scope and utility of our predictions. For example, lowering the threshold on gene body conservation to 10-15% makes it possible for the epigenome-based method to predict the expression status for approx. 90% of all genes in the target species, compared to approx. 80% for the orthology-based method. Given that both methods showed similar median accuracies of 75.1% and 79.9% (human reference) and of 71.5% and 78.1% (mouse reference), the epigenome-based provides a clear advantage over the orthology-based method by providing a substantially larger number of accurate predictions.

Finally, we evaluated the tradeoff between sensitivity and specificity in more detail for the epigenome-based method. We interpreted the class probabilities of its classifier as a measure of prediction confidence. We observed that the advantage of the epigenome-based method over the orthology-based method was strongest for relatively lenient thresholds (~0.5), at the cost of a slightly reduced accuracy (Figure 10; Additional file 1: Figure S6). For the most stringent thresholds (>0.9), the epigenome-based method continued to show a higher number of correct predictions than the orthology-based method, while the difference in accuracy between both methods approached zero. Based on the shape of the curves, a relatively stringent threshold of 0.75 is likely to constitute a suitable and broadly applicable tradeoff between sensitivity and specificity.

**Figure 10.**
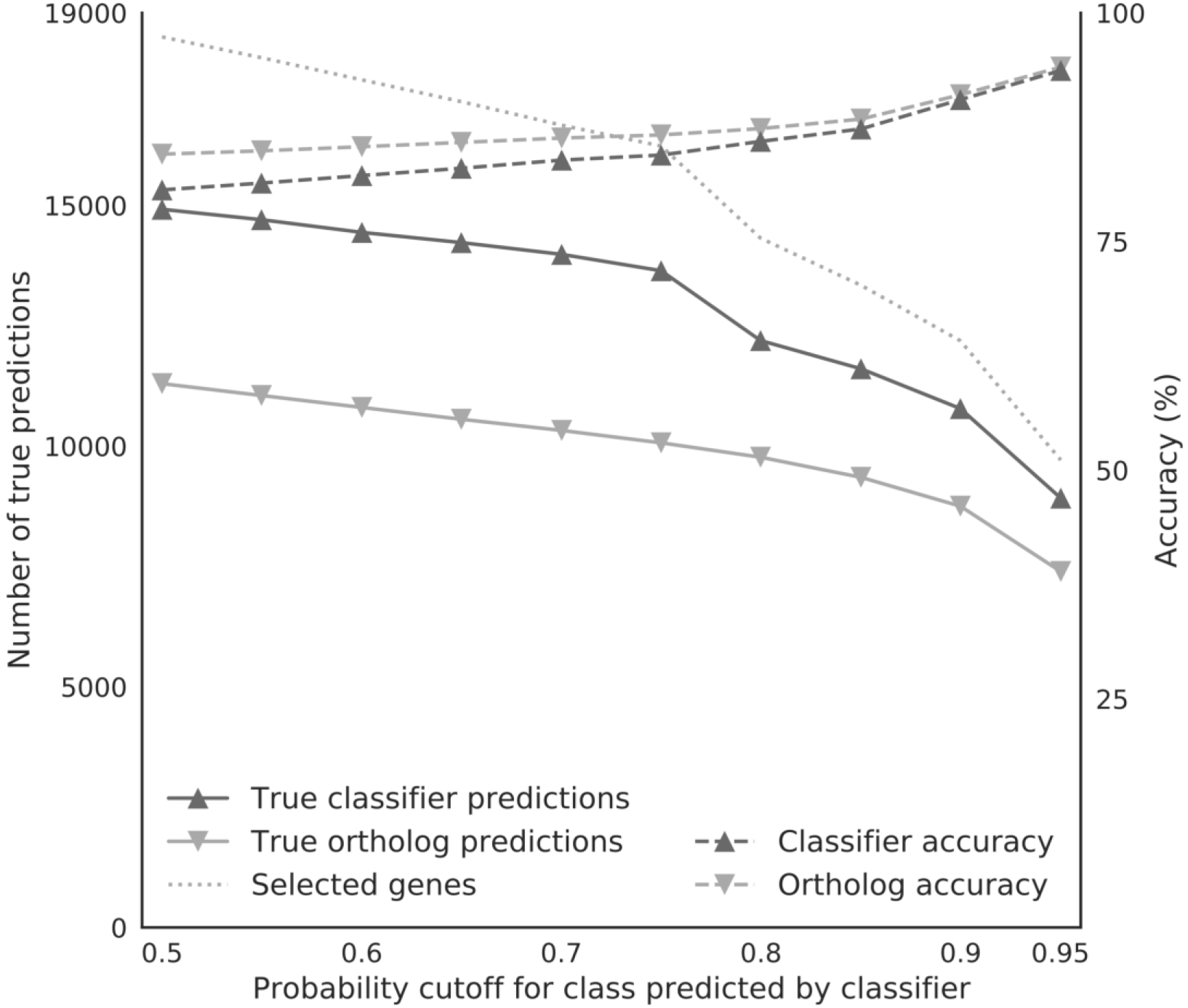
Effect of the classifier confidence threshold on prediction performance. Prediction performance of the epigenome-based approach (dark grey lines) and the orthology-based approach (light grey lines), as measured by the number of correct predictions (true positives and true negatives, left y-axis) and percent accuracy (right y-axis) for increasingly stringent cutoffs on the class probability estimates of the epigenome-based approach (x-axis, range from 0.5 to 0.95). Dotted line represent the number of genes selected at each threshold. Results are shown for human as reference and mouse as target species.

### Limitations of cross-species epigenome data transfer and prediction of gene expression

Despite these promising results using cross-species epigenome transfer and epigenome-based prediction of gene expression, the approach has certain limitations. Most importantly, whole-genome alignments cover only those genomic regions for which there is discernable conservation of the DNA sequence. For example, roughly 1 Gbp of DNA sequence were aligned between the human and mouse genome (90 million years of evolutionary distance), which corresponds to approximately a third of the human genome. At a threshold of at least 100 conserved base pairs for both the gene promoter and gene body, 92.2% (mouse) and 97.2% (human) of genes were amenable to cross-species transfer of reference epigenome data. These values remained high across larger evolutionary distances (Figure 11; Additional file 1: Figure S7), for example amounting to 98.8% in the comparison between human and opossum (160 million years) and 90.7% between human and chicken (312 million years).

**Figure 11.**
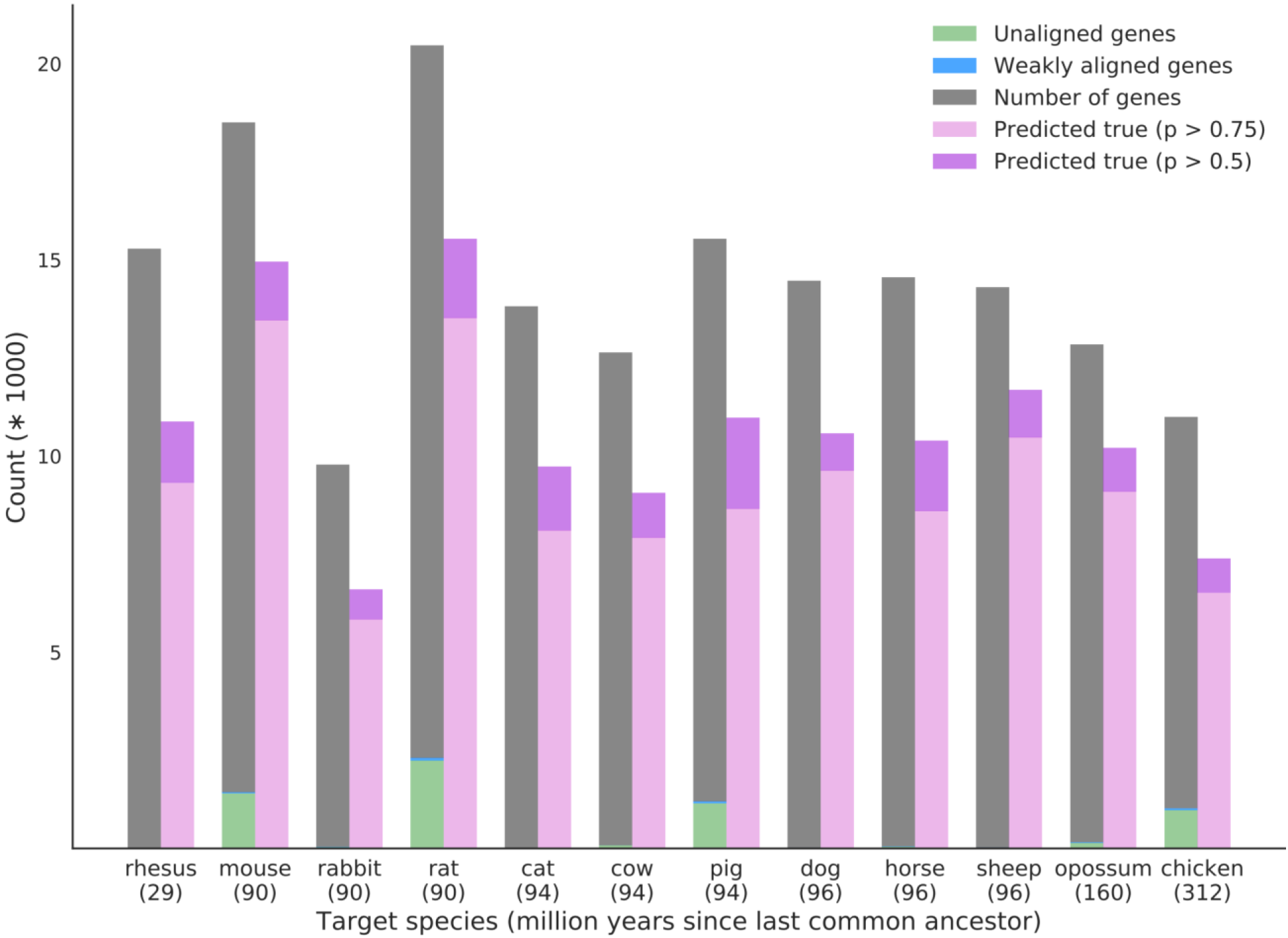
Epigenome-based prediction of gene expression for different target species. Bar plots showing for each species the total number of genes in the genome, the number of unaligned or weakly aligned genes relative to the reference species, and the average number of genes correctly predicted by the epigenome-based approach for two classifier confidence thresholds. Target species are sorted by evolutionary distance to the human reference along the x-axis (million years since last common ancestor indicated in parentheses).

We investigated the 487 human and 1,435 mouse genes that failed to meet our minimum alignment thresholds and which we therefore cannot predict using the epigenome-based method. Functionally characterization of these gene sets using the enrichR web service [34,35] identified an enrichment for Gene Ontology categories related to olfactory reception (Additional file 1: Figure S8A/B; Additional file 6), which is consistent with large species-specific differences in the repertoire of olfactory receptor genes between human and mouse [36]. We also analyzed the corresponding promoter regions for region set enrichment using the LOLA software [25], and we detected an enrichment of repetitive DNA elements (genomic duplications, satellite repeats, long terminal repeats, LINE repeats) as well as regions characterized by the heterochromatin mark H3K9me3 (Additional file 7). These results indicate that cross-species epigenome transfer is not well suited for analyzing species-specific gene families and repetitive heterochromatin regions.

Finally, we investigated whether the evolutionary age of individual genes may be associated with the accuracy of the epigenome-based predictions. Using a recently published dataset of gene age annotations [37], we found that genes whose expression status was predicted incorrectly tend to have a younger evolutionary age than those for which the expression status was predicted correctly (Figure 12; Additional file 1: Figure S9A/B). For example, the group of genes specific to mammals contained 52% more incorrectly predicted genes than would be expected based on the background distribution of gene ages; in contrast, the much larger group of genes shared across eukaryotes contained 7% more correctly predicted genes compared to expectation.

**Figure 12.**
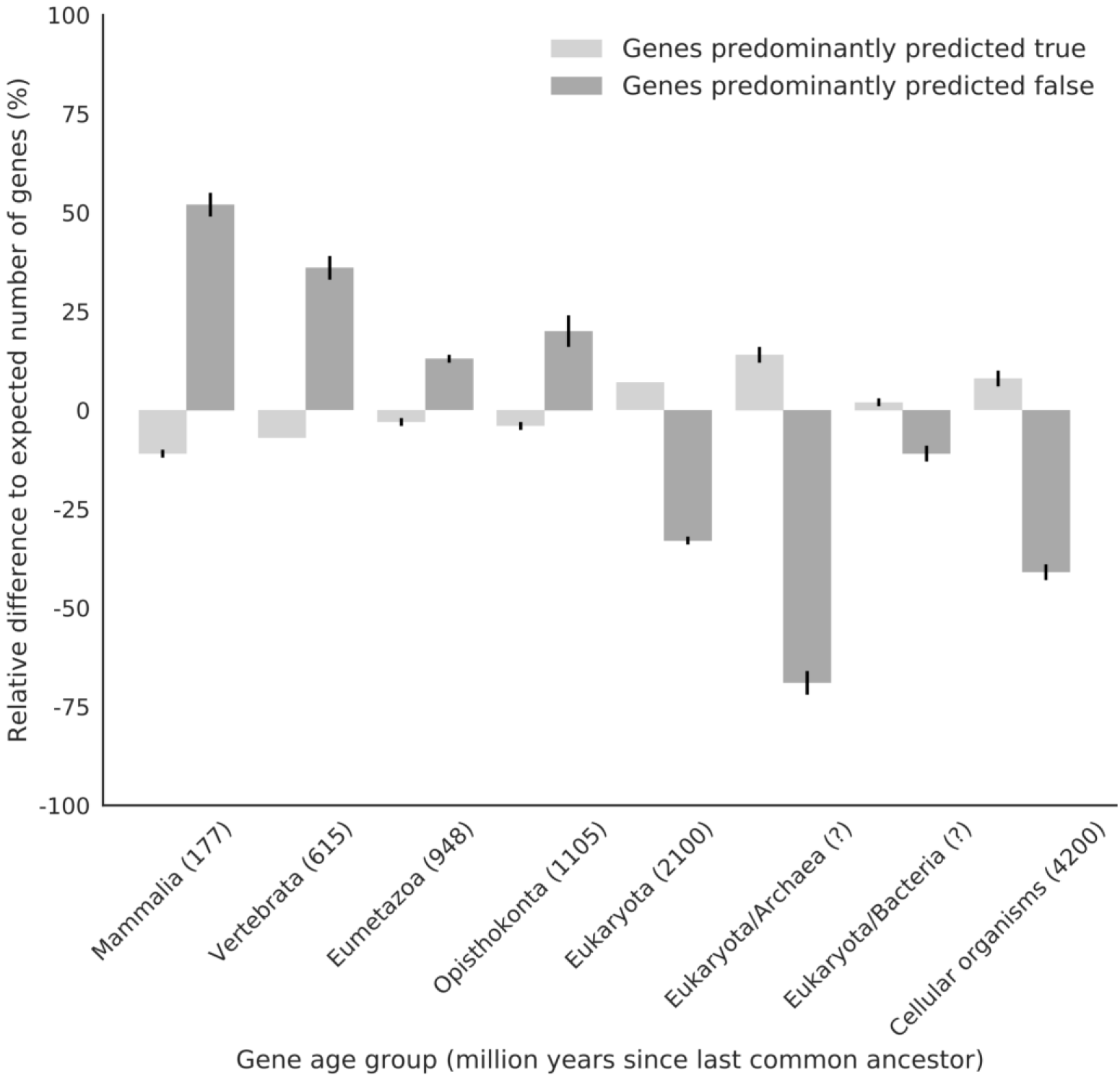
Association between gene age and epigenome-based prediction performance. Genes were stratified by their tendency to be predicted correctly or incorrectly by the epigenome-based approach and labeled with their annotated age. Bar heights indicate relative difference to the expected number of genes in each age group (percent values are shown for comparability across species). Error bars indicate +/- one standard deviation estimated by 1,000 bootstrap iterations. Numbers in parentheses indicate divergence time in million years relative to the family of Hominidae (great apes including human). Because of horizontal gene transfer, no divergence time estimates are given for the archaea and the bacteria group. The number of expected genes in each group is derived from the prior distribution for the respective gene age label.

## Discussion

Epigenome profiling is costly and labor-intensive, and comprehensive tissue-specific epigenome maps are currently available for only a few species. Here we explored cross-species extrapolation of epigenome data as a new approach that utilizes the large catalogs of human and mouse reference epigenomes to support research in non-model organisms.

We have developed a method for transferring epigenomes across species based on whole genome alignments, which we applied and validated in two complementary ways. First, we compared transferred and measured epigenomes for matched cell types between human and mouse (extensive epigenome profiles are available for validation in these two species), and we found that our method indeed retained the epigenome’s characteristic cell type specificity. Second, we combined cross-species epigenome transfer with epigenome-based prediction of gene expression, which adds an important dimension in the cross-species analysis of gene regulation and allowed us to validate our predictions in a larger number of target species for which transcriptome but no epigenome data were available. Again, we obtained high prediction accuracies, and our method accurately retained cell type specific regulatory patterns across species.

We observed broadly consistent results across twelve mammalian and one avian species included in our study, which span a spectrum of 20 million (mouse-rat) to 312 million (human-chicken and mouse-chicken) years of evolutionary distance. While it may be surprising that there was not a stronger deterioration of prediction performance as evolutionary distances increased, all of the species included in our study share a highly conserved epigenetic machinery, and their genomes are sufficiently conserved that it is still possible to identify cross-species homology in gene promoters and gene bodies using whole genome alignments. Moreover, we found that evolutionary old genes were predicted with higher accuracy than more recently evolved genes, which is likely to contribute to the robustness of predictions across a wide evolutionary range.

Our method does not have major limitations that would hinder its broad application, except that it requires a reference genome for all included species. The more complete and more accurate the reference genome assemblies of the target species, the higher will be the quality of whole genome alignments, and – as a direct consequence – the performance of our method. Nevertheless, because our method transfers epigenome data between locally aligned regions, it is not restricted to high-quality genomes and can be expected to cope well with highly fragmented genome assemblies, thus facilitating its application to understudied non-model organisms.

## Conclusion

Our results show that cross-species transfer of epigenome data is possible among mammalian (and avian) species, and that the transferred epigenomes not only retain tissue specificity but also enable tissue-specific prediction of gene expression. We can thus conclude that the tissue-specific links between epigenome profiles and gene expression are well conserved across the analyzed species. Cross-species epigenome transfer and prediction can help address the current dearth of epigenome data for non-model organisms. Importantly, computational prediction is not meant to replace experimental analysis, but to complement it by providing access to a larger number of species and more tissues within each species. It will also be interesting to compare predicted epigenome data with experimentally measured epigenome data, in order to find genomic regions in which the measurement deviates from the prediction. Such regions would be strong candidates for species-specific epigenome regulation and promising targets for in-depth biological investigation. We thus conclude that bioinformatic approaches for cross-species epigenome mapping can reasonably complement well-established experimental methods.

## Materials and Methods

### Included species, genome assemblies, and gene models

We included twelve mammalian species (human [hg19], rhesus [rheMac2], mouse [mm9], rat [rn5], rabbit [oryCun2], pig [susScr2], cow [bosTau7], sheep [oviAri3], horse [equCab2], dog [canFam3], cat [felCat5], opossum [monDom5]) and one avian species (chicken [galGal3]), based on the following selection criteria: (i) complete genome assemblies and whole-genome alignments with at least one of the two reference species (human, mouse) were available from the UCSC Genome Browser [39]; (ii) gene models for the relevant assemblies were available from one of three sources (GENCODE [40]: human v19, mouse vM1; The Bovine Genome Database [41]: cow Ensembl75; UCSC Genome Browser tables ensGene, ensemblSource and ensemblToGeneName: all other species); (iii) epigenome profiles including histone H3K4me3, H3K27ac, and H3K36me3 as well as transcriptome data were available for defined tissues / cell types in the reference species (Additional file 2); and (iv) transcriptome data were available for at least some of these tissues / cell types in the target species. An overview of the evolutionary relationships for all species included in this study is provided as a phylogenetic tree generated using the Time-Tree service [42] (Additional file 1: Figure S1). To alleviate the effect of differences in annotation quality, all gene models were reduced to protein-coding transcripts/genes. Additionally, only transcripts tagged as “Consensus CDS (CCDS)” were selected in the GENCODE annotations. All analyses were restricted to genes located on the autosomes, excluding the sex chromosomes.

Promoter regions were defined as 1.5 kilobase windows around the transcription start site (−1,000 bp to +500 bp), and genes with a gene body length of less than 750 bp were discarded.

### Whole-genome alignments, gene orthologs, and evolutionary conservation

Whole-genome alignments between the reference species (human, mouse) and target species were downloaded from the UCSC Genome Browser in the form of chain files [43] and processed as described in the UCSC Genome Wiki to derive reciprocal best chains [44]. The reciprocal best chains were further processed using CrossMap [45] and custom scripts to build pairwise symmetric alignment blocks. Genes with less than 100 aligned bases in their promoter and in their body were considered weakly aligned. Information on gene orthologs was downloaded from OrthoDB [33], and lists of 1-to-1 orthologs for each pair of species and, separately, for all 13 species combined were extracted from the annotation data using custom scripts.

### Epigenome and transcriptome data preprocessing

Publicly available reference epigenomes for the reference species (human, mouse) were obtained from ENCODE, DEEP, and BLUEPRINT projects (Additional file 2). The resulting dataset included three histone marks (H3K4me3, H3K27ac, H3K36me3), three cell types (embryonic stem cells, naïve CD4+ T cells and hepatocytes), and a total of nine epigenome profiles for human, as well as three histone marks (H3K4me3, H3K27ac, H3K36me3), five cell/tissue types (embryonic stem cells, naïve CD4+ T cells, whole liver, kidney and heart), and a total of 17 epigenome profiles for mouse. Tissue-specific transcriptome profiles were obtained from ENCODE, DEEP, and public repositories (SRA/ENA). Where available, epigenome profiles were downloaded in form of histone signal tracks (bigWig format). For the BLUEPRINT mouse data, which were not available in preprocessed form, reads were mapped using bowtie2 v2.3.3.1 with the preset *--sensitive*, and signal tracks were generated using bamCoverage v2.5.3 from the deepTools software suite [46]. To prepare the epigenome profiles for the analysis, biological replicates from the same laboratory were merged by taking the mean. All resulting epigenome signal tracks were quantile normalized per project and clipped at the 99.95 percentile to alleviate the effect of outliers. Transcriptome data were processed with Salmon v0.8.2 using the following parameters: *--forgettingFactor 0.8 --useVBOpt --seqBias --gcBias --geneMap*, aggregating transcript-level abundance estimates into gene-level estimates. Finally, the gene expression values were subjected to quantile normalization, resulting in transcript per million (TPM) values that were used for further analysis.

### Epigenome-based prediction of gene expression

All prediction models were implemented in Python3, using libraries from the SciPy ecosystem for scientific computing [47,48]. Histone signal tracks were masked to exclude non-conserved regions according to pairwise genome alignments between the reference and target species. Prediction attributes were derived from these masked signal tracks by averaging the signal across each gene promoter (H3K4me3, H3K27ac) and gene body (H3K36me3). Gradient boosting classifiers from the scikit-learn library [49] were trained using these histone signal intensities as prediction attributes and the gene expression status (on/high: TPM ≥ 1; off/low: TPM < 1) as target variable (this binary thresholding strategy was motivated by previous studies [28, 50–52]). Each training dataset was randomly subsampled to balance class frequencies, and model hyperparameters were tuned using 5-fold cross-validation on this subsampled training dataset. The best model according to the results of the cross-validation was refit using the full set of training data.

### Genomic region enrichment analysis

Region sets were analyzed for significant enrichment using the LOLA software [25]. For the human genome, the LOLA Core region set was used. In addition, we created a custom region set for both the human and mouse genome, comprising various sets of transcription factor binding sites as well as histone modification peaks from the DeepBlue repository [53]. For each LOLA analysis, we filtered the results and retained enriched region sets only if the support (i.e., number of regions covered) was at least 5 and the multiple-testing corrected statistical significance (q-value) was below 0.05. We manually selected 10-15 entries from the top ranking region sets for visualization and provide the full list of LOLA enrichments in Additional files 5 and 7.

### Gene age annotation

To evaluate the evolutionary age of each gene, we obtained gene age annotations for the following species from a recent publication [37]: chicken, cow, dog, human, mouse, opossum, rat, and rhesus macaque. UniProt identifiers were mapped to Ensembl gene identifiers using a web-based service [54]. For visualization, we selected all species for which at least 80% of the identifiers could be mapped (human 97%; mouse 83%; rhesus 81%) and plotted the observed gene age distribution relative to the expected distribution based on the priors for the different gene age labels (Figure 12, Additional file 1: Figure S9A/B).

## Declarations

### Availability of data and material

All datasets used are listed in Additional file 2 and publicly available. Transferred epigenome are provided in Additional files 3 and 4. Program code and instructions to replicate all results are openly available in a github repository (https://doi.org/10.17617/1.69).

### Competing interests

None declared.

### Funding

This work was performed in the context of the German Epigenome Project (DEEP, German Science Ministry grant no. 01KU1216A) and the BLUEPRINT project (European Union’s Seventh Framework Programme grant 282510). C.B. is supported by a New Frontiers Group award of the Austrian Academy of Sciences and by an ERC Starting Grant (European Union’s Horizon 2020 research and innovation programme, grant agreement no. 679146).

### Authors’ contributions

P.E. and C.B. conceptualized the project with input from T.L.; P.E. analyzed the data and wrote the first draft of the manuscript; all authors contributed to the writing of the manuscript.

## Acknowledgements

We acknowledge the ENCODE, DEEP, and BLUEPRINT consortia for providing epigenome data for the reference species. Additionally, we thank Anna Hake (née Feldmann), Prabhav Kalaghatgi, Johanna Klughammer, and Nora Speicher for helpful discussions.

## Additional files

Name: Additional file 1

Format: doc text document (MS Word / LibreOffice Writer)

Title: Supplementary figures

Description: Supplementary figures

Name: Additional file 2

Format: tsv tab-separated values (MS Excel / LibreOffice Calc)

Title: Datasets overview

Description: Summary of epigenome and transcriptome datasets used in this study

Name: Additional file 3

Format: tsv tab-separated values (MS Excel / LibreOffice Calc)

Title: Epigenome predictions (human reference)

Description: Epigenome data for gene promoters and gene bodies transferred from human (as the reference species) to twelve target species

Name: Additional file 4

Format: tsv tab-separated values (MS Excel / LibreOffice Calc)

Title: Epigenome predictions (mouse reference)

Description: Epigenome data for gene promoters and gene bodies transferred from mouse (as the reference species) to twelve target species

Name: Additional file 5

Format: zip compressed tab-separated values (7-Zip, MS Excel / LibreOffice Calc)

Title: LOLA results for transferred epigenome data

Description: Region set enrichment analysis using LOLA for tissue-specific epigenome transfer between human and mouse (and vice versa)

Name: Additional file 6

Format: zip compressed tab-separated values (7-Zip, MS Excel / LibreOffice Calc)

Title: enrichR results for weakly aligned genes

Description: Gene set enrichment analysis using enrichR for weakly aligned genes in human and mouse

Name: Additional file 7

Format: zip compressed tab-separated values

Title: LOLA results for weakly aligned genes

Description: Region set enrichment analysis using LOLA for weakly aligned genes in human and mouse

## References

1. Boffelli D, Nobrega MA, Rubin EM. Comparative genomics at the vertebrate extremes. Nat Rev Genet. 2004;5:456–65.

2. Breschi A, Gingeras TR, Guigó R. Comparative transcriptomics in human and mouse. Nat Rev Genet. 2017.

3. Koonin E.V. Are There Laws of Genome Evolution? Bourne PE, editor. PLoS Comput Biol. 2011;7:e1002173.

4. Lehner B. Genotype to phenotype: lessons from model organisms for human genetics. Nat Rev Genet. 2013;14:168–78.

5. Meadows JRS, Lindblad-Toh K. Dissecting evolution and disease using comparative vertebrate genomics. Nat Rev Genet. 2017;18:624–36.

6. Villar D, Flicek P, Odom DT. Evolution of transcription factor binding in metazoans - mechanisms and functional implications. Nat Rev Genet. 2014;15:221–33.

7. Wilson MD, Odom DT. Evolution of transcriptional control in mammals. Curr Opin Genet Dev. 2009;19:579–85.

8. IHEC Consortium. International Human Epigenome Consortium [Internet]. 2010. Available from: http://ihec-epigenomes.org

9. Adams D, Altucci L, Antonarakis S, Ballesteros J, Beck S, Bird A, et al. BLUEPRINT to decode the epigenetic signature written in blood. Nat Biotechnol. 2012;30.

10. DEEP Consortium. Deutsches Epigenom Programm [Internet]. 2012. Available from: www.deutsches-epigenom-programm.de

11. The ENCODE Project Consortium, Dunham I, Kundaje A, Aldred SF, Collins PJ, et al. An integrated encyclopedia of DNA elements in the human genome. Nature. 2012;489:57–74.

12. Stamatoyannopoulos JA, Snyder M, Hardison R, Ren B, Gingeras T, Gilbert DM, et al. An encyclopedia of mouse DNA elements (Mouse ENCODE). Genome Biol. 2012;13:418.

13. Heintzman ND, Hon GC, Hawkins RD, Kheradpour P, Stark A, Harp LF, et al. Histone modifications at human enhancers reflect global cell-type-specific gene expression. Nature. 2009;459:108–12.

14. Gutierrez-Arcelus M, Ongen H, Lappalainen T, Montgomery SB, Buil A, Yurovsky A, et al. Tissue-Specific Effects of Genetic and Epigenetic Variation on Gene Regulation and Splicing. PLOS Genet. 2015;11:e1004958.

15. Thurman RE, Rynes E, Humbert R, Vierstra J, Maurano MT, Haugen E, et al. The accessible chromatin landscape of the human genome. Nature. 2012;489:75–82.

16. Villar D, Berthelot C, Aldridge S, Rayner TF, Lukk M, Pignatelli M, et al. Enhancer Evolution across 20 Mammalian Species. Cell. 2015;160:554–66.

17. Yang S, Oksenberg N, Takayama S, Heo S-J, Poliakov A, Ahituv N, et al. Functionally conserved enhancers with divergent sequences in distant vertebrates. BMC Genomics. 2015;16:882.

18. Ellegren H. Genome sequencing and population genomics in non-model organisms. Trends Ecol Evol. 2014;29:51–63.

19. Koepfli K-P, Paten B, O’Brien SJ. The Genome 10K Project: A Way Forward. Annu Rev Anim Biosci. 2015;3:57–111.

20. Schmidt D, Wilson MD, Ballester B, Schwalie PC, Brown GD, Marshall A, et al. Five-Vertebrate ChIP-seq Reveals the Evolutionary Dynamics of Transcription Factor Binding. Science 2010;328:1036–40.

21. Long HK, Sims D, Heger A, Blackledge NP, Kutter C, Wright ML, et al. Epigenetic conservation at gene regulatory elements revealed by non-methylated DNA profiling in seven vertebrates. Elife. 2013;2:e00348.

22. The Genome 10K Project. Genome 10K: a proposal to obtain whole-genome sequence for 10,000 vertebrate species. J Hered. 2009;100:659–74.

23. Yue F, Cheng Y, Breschi A, Vierstra J, Wu W, Ryba T, et al. A comparative encyclopedia of DNA elements in the mouse genome. Nature. 2014;515:355–64.

24. Durek P, Nordström K, Gasparoni G, Salhab A, Kressler C, de Almeida M, et al. Epigenomic Profiling of Human CD4 + T Cells Supports a Linear Differentiation Model and Highlights Molecular Regulators of Memory Development. Immunity. 2016;45:1148–61.

25. Sheffield NC, Bock C. LOLA: enrichment analysis for genomic region sets and regulatory elements in R and Bioconductor. Bioinformatics. 2016;32:587–9.

26. Birney E, Stamatoyannopoulos JA, Dutta A, Guigó R, Gingeras TR, Margulies EH, et al. Identification and analysis of functional elements in 1% of the human genome by the ENCODE pilot project. Nature. 2007;447:799–816.

27. Karlić R, Chung H-R, Lasserre J, Vlahovicek K, Vingron M. Histone modification levels are predictive for gene expression. Proc Natl Acad Sci U S A. 2010;107:2926–31.

28. Singh R, Lanchantin J, Robins G, Qi Y. DeepChrome: deep-learning for predicting gene expression from histone modifications. Bioinformatics. 2016;32:i639–48.

29. Clever D, Roychoudhuri R, Constantinides MG, Askenase MH, Sukumar M, Klebanoff CA, et al. Oxygen Sensing by T Cells Establishes an Immunologically Tolerant Metastatic Niche. Cell. 2016;166:1117–1131.e14.

30. Fushan AA, Turanov AA, Lee S-G, Kim EB, Lobanov A.V., Yim SH, et al. Gene expression defines natural changes in mammalian lifespan. Aging Cell. 2015;14:352–65.

31. Merkin J, Russell C, Chen P, Burge CB. Evolutionary Dynamics of Gene and Isoform Regulation in Mammalian Tissues. Science. 2012;338:1593–9.

32. Jiang Y, Xie M, Chen W, Talbot R, Maddox JF, Faraut T, et al. The sheep genome illuminates biology of the rumen and lipid metabolism. Science. 2014;344:1168–73.

33. Kriventseva E V, Tegenfeldt F, Petty TJ, Waterhouse RM, Simao FA, Pozdnyakov IA, et al. OrthoDB v8: update of the hierarchical catalog of orthologs and the underlying free software. Nucleic Acids Res. 2015;43:D250–6.

34. Chen EY, Tan CM, Kou Y, Duan Q, Wang Z, Meirelles G, et al. Enrichr: interactive and collaborative HTML5 gene list enrichment analysis tool. BMC Bioinformatics. 2013;14:128.

35. Kuleshov M V., Jones MR, Rouillard AD, Fernandez NF, Duan Q, Wang Z, et al. Enrichr: a comprehensive gene set enrichment analysis web server 2016 update. Nucleic Acids Res. 2016;44:W90–7.

36. Niimura Y, Nei M. Comparative evolutionary analysis of olfactory receptor gene clusters between humans and mice. Gene. 2005;346:13–21.

37. Liebeskind BJ, McWhite CD, Marcotte EM. Towards Consensus Gene Ages. Genome Biol Evol. 2016;8:1812–23.

38. Brawand D, Soumillon M, Necsulea A, Julien P, Csárdi G, Harrigan P, et al. The evolution of gene expression levels in mammalian organs. Nature. 2011;478:343–8.

39. Zhi H, Li X, Wang P, Gao Y, Gao B, Zhou D, et al. The UCSC Genome Browser database: 2018 update. Nucleic Acids Res. 2017;46.

40. Harrow J, Frankish A, Gonzalez JM, Tapanari E, Diekhans M, Kokocinski F, et al. GENCODE: the reference human genome annotation for The ENCODE Project. Genome Res. 2012;22:1760–74.

41. Elsik CG, Unni DR, Diesh CM, Tayal A, Emery ML, Nguyen HN, et al. Bovine Genome Database: new tools for gleaning function from the Bos taurus genome. Nucleic Acids Res. 2016;44:D834–9.

42. Hedges SB, Marin J, Suleski M, Paymer M, Kumar S. Tree of Life Reveals Clock-Like Speciation and Diversification. Mol Biol Evol. 2015;32:835–45.

43. Kent WJ, Baertsch R, Hinrichs A, Miller W, Haussler D. Evolution’s cauldron: duplication, deletion, and rearrangement in the mouse and human genomes. Proc Natl Acad Sci U S A. 2003;100:11484–9.

44. UCSC Genome Browser. UCSC Genome Wiki: reciprocal chain/nets [Internet]. 2016. Available from: http://genomewiki.ucsc.edu/index.php/HowTo:_Syntenic_Net_or_Reciprocal_Best

45. Zhao H, Sun Z, Wang J, Huang H, Kocher J-P, Wang L. CrossMap: a versatile tool for coordinate conversion between genome assemblies. Bioinformatics. 2014;30:1006–7.

46. Ramírez F, Dündar F, Diehl S, Grüning B a, Manke T. deepTools: a flexible platform for exploring deep-sequencing data. Nucleic Acids Res. 2014;42:187–91.

47. Millman KJ, Aivazis M. Python for Scientists and Engineers. Comput Sci Eng. 2011;13:9–12.

48. Oliphant TE. Python for Scientific Computing. Comput Sci Eng. 2007;9:10–20.

49. Pedregosa F, Varoquaux G, Gramfort A, Michel V, Thirion B, Grisel O, et al. Scikit-learn: Machine Learning in Python. J Mach Learn Res. 2011;12:2825–30.

50. Budden DM, Hurley DG, Cursons J, Markham JF, Davis MJ, Crampin EJ. Predicting expression: the complementary power of histone modification and transcription factor binding data. Epigenetics Chromatin. 2014;7:36.

51. Cheng C, Yan K-K, Yip KY, Rozowsky J, Alexander R, Shou C, et al. A statistical framework for modeling gene expression using chromatin features and application to modENCODE datasets. Genome Biol. 2011;12:R15.

52. Dong X, Greven MC, Kundaje A, Djebali S, Brown JB, Cheng C, et al. Modeling gene expression using chromatin features in various cellular contexts. Genome Biol. 2012;13:R53.

53. Albrecht F, List M, Bock C, Lengauer T. DeepBlue epigenomic data server: programmatic data retrieval and analysis of epigenome region sets. Nucleic Acids Res. 2016;44:W581–6.

54. UniProt. UniProt ID mapping [Internet]. Available from: www.uniprot.org/uploadlists

